# Recommendations for Interpreting the Loss of Function PVS1 ACMG/AMP Variant Criteria

**DOI:** 10.1101/313718

**Authors:** Ahmad N. Abou Tayoun, Tina Pesaran, Marina T. DiStefano, Andrea Oza, Heidi L. Rehm, Leslie G. Biesecker, Steven M. Harrison, on behalf of the ClinGen Sequence Variant Interpretation Working Group

## Abstract

The 2015 ACMG/AMP sequence variant interpretation guideline provided a framework for classifying variants based on several benign and pathogenic evidence criteria. This guideline includes a pathogenic criterion (PVS1) for predicted loss of function variants. However, the guideline did not elaborate on the specific considerations for the different types of loss of function variants, nor did it provide decision-making pathways assimilating information about the variant type, its location within the gene, or any additional evidence for the likelihood of a true null effect. Furthermore, the ACMG/AMP guideline did not take into account the relative strengths for each evidence type and the final outcome of their combinations with respect to PVS1 strength. Finally, criteria specifying the genes for which PVS1 can be used are still missing. Here, as part of the Clinical Genomic Resource (ClinGen) Sequence Variant Interpretation (SVI) Working Group’s goal of refining ACMG/AMP criteria, we provide recommendations for applying the PVS1 rule using detailed guidance addressing all the above-mentioned gaps. We evaluate the performance of the refined rule using heterogeneous types of loss of function variants (n = 56) curated by seven disease-specific groups across ten genes. Our recommendations will facilitate consistent and accurate interpretation of predicted loss of function variants.

**GRANT NUMBERS:** Research reported in this publication was supported by the National Human Genome Research Institute (NHGRI) under award number U41HG006834. LGB was supported by the Intramural Research Program of the NHGRI grant number HG200359 09.

## INTRODUCTION

In 2015, the American College of Medical Genetics and Genomics (ACMG) and the Association for Molecular Pathology (AMP) published a joint guideline that provides a framework for sequence variant interpretation (Richards et al., 2015). The guideline defined 28 criteria, each with an assigned code, that addressed distinct types of variant evidence. Each criterion code was assigned a direction, benign (B) or pathogenic (P), and a level of strength: stand-alone (A), very strong (VS), strong (S), moderate (M), or supporting (P). Combining rules for these criteria were also proposed to determine the pathogenicity of sequence variants.

The only criterion designated with Very Strong strength level for pathogenicity in the ACMG/AMP guideline was PVS1 which was defined as “null variant (nonsense, frameshift, canonical ±1 or 2 splice sites, initiation codon, single or multi-exon deletion) in a gene where loss-of-function (LoF) is a known mechanism of disease” (Richards et al., 2015). A combination of this rule and only one moderate or two supporting pathogenicity criteria lead to a likely pathogenic or pathogenic classification, respectively in the original ACMG/AMP recommendations. Given the weighting of this criterion as very strong and the consequent impact of any potential inappropriate usage, detailed guidance on its application is critical. Despite addressing general considerations associated with PVS1 usage including disease mechanism, splice variant effects, nonsense-mediated decay (NMD), and alternative splicing, the ACMG/AMP guideline did not provide guidance for how to account for these considerations during variant assessment and determination of whether PVS1 was applicable. Additionally, while the ACMG/AMP guideline stated that criteria listed as one strength can be moved to another strength using professional judgment, no guidance was provided regarding instances in which the strength of PVS1 should be decreased to Strong (PVS1_Strong), Moderate (PVS1_Moderate), or Supporting (PVS1_Supporting) strength level.

The NIH-funded Clinical Genome Resource (ClinGen) established the Sequence Variant Interpretation (SVI) working group (https://www.clinicalgenome.org/working-groups/sequence-variant-interpretation/) to refine and evolve the ACMG/AMG rules for accurate and consistent clinical application, as well as harmonize disease-focused specification of the guidelines by Expert Panels (Gelb et al., 2018; Kelly et al., 2018).

In this report, we provide detailed recommendations by the SVI working group for interpretation of the PVS1 rule. These recommendations provide criteria for determining if LoF is a disease mechanism for the associated gene/disease and address variant type-specific considerations (nonsense, frameshift, initiation codon, invariant splice site, deletion and duplication) in the context of gene structure and pathophysiologic mechanisms, such as NMD or alternative splicing. In addition, we assign varying modifications of PVS1 strength based on assimilation of the available evidence (“Guidance on how to rename criteria codes when strength of evidence is modified” section on SVI webpage). Finally, 56 LoF variants of varying variant type and across multiple genes were curated by ClinGen disease-specific working groups to determine if the recommendations were easy to follow, accounted for all LoF scenarios encountered, and if the working group agreed with the specified PVS1 strength level for each tested variant.

## METHODS

In July 2017, the SVI Working Group, representing clinical geneticists, genetic counselors, genomic researchers, and clinical laboratory geneticists, held a two-day in-person meeting in Boston to specifically refine and extend several ACMG/AMP criteria including the PVS1 rule.

During this meeting, the group outlined a detailed framework for evolving the previous PVS1 rule into the current recommendations in this report. Subsequently, a smaller group within the ClinGen Hearing Loss (HL) Working Group continued further refinement of this rule through weekly conference calls and solicited feedback from the SVI Working group via monthly conference calls.

In October 2017, the SVI Working Group held a second in-person meeting at the American Society of Human Genetics (ASHG) meeting in Orlando. During that meeting, the group finalized a first recommendation draft and provided comments for additional refinements that were addressed through the HL group and later approved by the SVI Working Group.

Throughout the PVS1 rule refinement process, we used expert opinions, empirical data in the literature, and unpublished observations from participating research and clinical laboratories. In addition, to ensure comprehensive utility of the new rule, seven ClinGen Clinical Domain Working Groups (CDWGs) were asked to use this rule to classify five to ten LoF variants each in their genes of interest (total 56 variants in ten genes). Their feedback was then incorporated into the final PVS1 recommendations.

## RESULTS

### RECOMMENDATION FOR APPLICATION OF PVS1 CRITERION

The SVI working group has created a PVS1 decision tree (**Figure 1**) to guide curators on the applicable PVS1 strength level depending on variant type (duplication, deletion, splice site, nonsense/frameshift, initiation codon) and variant features (such as predicted impact, location in the gene, and inclusion of impacted exon). The current decision tree format assumes that the gene/disease association is at a Strong or Definitive clinical validity level (Strande et al., 2017) and that LoF is an established disease mechanism (see “Disease Mechanism” section).

**Figure 1.**
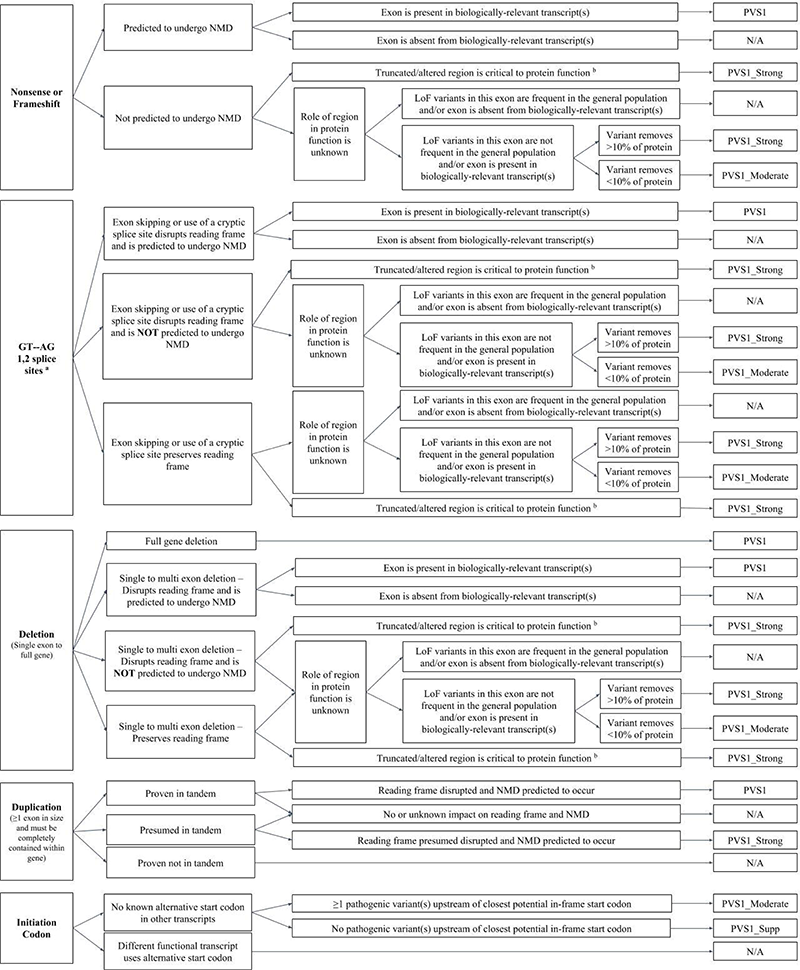
PVS1 decision tree. Refer to text for detailed description. NMD, nonsense-mediated decay; LoF, loss of function. a, This criterion should not be applied in combination with in silico splicing predictions (PP3). Additionally, splice site variants must have no detectable nearby (+/− 20 nts) strong consensus splice sequence that may reconstitute in-frame splicing. b, Relevant domain indicated by experimental evidence proving a critical role of the domain and/or presence of non-truncating pathogenic variants in the region.

#### PVS1 Strength Levels

The SVI Working Group has recently modeled the ACMG/AMP variant classification guidelines into a Bayesian framework whereby the relative odds of pathogenicity for supporting, moderate, strong, and very strong pathogenic evidence were estimated to be 2.08, 4.33, 18.7, and 350, respectively (Tavtigian et al., 2018). In refining the PVS1 criteria, the Working Group determined that not all putative LoF variants have equal strengths, and that the PVS1 strength level can vary depending on the available evidence for each variant type. Therefore, we divided this criterion into PVS1, PVS1_Strong, PVS1_Moderate, and PVS1_Supporting. Although we did not quantify each evidence type or a combination thereof, to maintain consistency in interpreting this rule, the above relative odds of pathogenicity were considered before assigning a PVS1 strength level. Lastly, at the Moderate strength level, there is potential overlap in usage of PVS1_Moderate and PM4 (protein length changing variant). To prevent double-counting of this evidence type, we recommend that PM4 should not be applied for any variant in which PVS1, at any strength level, is also applied.

#### Alternate transcripts and nonsense mediated mRNA decay (NMD) considerations

The predicted impact of a premature termination codon on an mRNA and/or a protein product depends on the location of the new termination codon within the most biologically relevant transcript(s). Generally, NMD is not predicted to occur if the premature termination codon occurs in the 3’ most exon or within the 3’-most 50 nucleotides of the penultimate exon (Chang, Imam, & Wilkinson, 2007; Lewis, Green, & Brenner, 2003). When NMD is not predicted to occur, it is important to determine if the truncated or altered region is critical to protein function, often indicated by experimental or clinical evidence – such as pathogenic variants downstream of the new stop codon – supporting the biological relevance of the C-terminal region. With this evidence, we estimated the likelihood of pathogenicity to mount to at least ~19:1 odds of pathogenicity consistent with PVS1_Strong assignment (**Figure 1**). If there was no variant or functional evidence indicating the truncated region is critical to protein function then assessing tolerance of the exon to LoF variants and inclusion in biologically-relevant transcripts can be helpful. In this case, if the affected exon was neither enriched with high frequency LoF variants in the general population or absent from biologically-relevant transcripts (either of which would inactivate PVS1 usage, see below), then the length of the missing region factors into PVS1 strength level decision making. In this scenario, and in the absence of pertinent data, the SVI Working Group reached a consensus agreement that removing >10% of the protein product is more likely to have a loss of function effect (PVS1_Strong) compared to variants that remove <10% of the protein (PVS1_Moderate) (**Figure 1**). We acknowledge that empirical data are needed to support and further refine this generic rule and we anticipate that disease-specific groups will specify based on expert knowledge of their genes of interest.

If the putative LoF variant occurs in an exon upstream of where NMD is predicted to occur, then alternative splicing of this exon from the major or most biologically relevant transcript must be assessed before application of PVS1 (**Figure 1**). Generally, a transcript or exon is considered biologically relevant based on functional and/or expression evidence. In addition, presence of pathogenic variants, in an exon is supportive of inclusion of the exon in the biologically relevant transcript.

In general, PVS1 at any strength level should not be applied if the putative loss of function variant affects exon(s) which is/are missing from alternate biologically relevant transcript(s) **OR** is/are enriched for high frequency LoF variants in the general population (**Figure 1**). The frequency threshold at which a LoF variant in the general population is considered to be high is dependent on specific gene and/or disease attributes such as prevalence, gene contribution, allelic heterogeneity, mode of inheritance and penetrance. Each disease group should determine such cutoffs in the process of estimating their allele frequency thresholds.

#### Variant Type Considerations

##### Nonsense and frameshift variants

The first step in the interpretation process for nonsense and frameshift variants includes determination of the location of the new termination codon within the most biologically relevant transcript. As explained above, this is critical to determining if NMD is predicted to occur, or if the putative LoF variant is in a non-essential exon that is either alternatively spliced from the major transcript, enriched with high frequency LoF variants in the general population, and/or removes a downstream region that is not critical to protein function. Different combinations of these variables will lead to different outcomes with respect to using PVS1, at any strength level, or not at all as illustrated in **Figure 1**.

##### Canonical ±1,2 splice variants

Mutations of the canonical ± 1 or 2 splice sites are often presumed to have loss of function effects. The major consensus nucleotides at the U2 spliceosome donor and acceptor splice nucleotides are GT and AG, respectively. It is important to note that for those variants, the PP3 (*in silico* splicing prediction) criterion should not be used to avoid double counting the same predictive evidence used to assign PVS1. When interpreting ± 1,2 splice variants, it is useful to predict the impact that altered splicing may have on the protein reading frame (**Figure 1**). While it is challenging to predict the effects of splice site variants (e.g., skipped exon, use of a cryptic splice site, etc.) without RNA studies, it is useful to search for cryptic splice sites as well as anticipate the impact of exon skipping or cryptic splice site usage. First, one should assess nearby (±20bp) sequences for any cryptic splice sites that may reconstitute in-frame splicing. Next, one should determine if the nucleotide sequence of the exon is divisible by three and therefore could lead to an in-frame deletion in an otherwise intact transcript or if it is not divisible by three and would predict a frameshift if the exon is simply skipped. Then the consequence of use of any cryptic splice site as well as exon skipping should be assessed and the lowest strength of PVS1 should be applied among the scenarios. Similar approaches to assessing NMD, the biological relevance of the exon and protein region, as described above for nonsense and frameshift variants, should then be applied (**Figure 1**).

##### Initiation codon variants

Functional studies have shown that start re-initiation can be very robust occurring at alternate ATG or non-ATG sites downstream and even upstream of the lost original start site (Bazykin & Kochetov, 2011; Drabkin & RajBhandary, 1998; Lee et al., 2012; Na et al., 2018; Starck et al., 2012; Wan & Qian, 2014; Zur & Tuller, 2013). Based on these findings, the SVI Working Group does not generally recommend assigning PVS1 or PVS1_Strong for start loss variants. If alternative functional gene transcripts (i.e., found in transcript or expression databases) use an alternative start codon, then we recommend not applying PVS1 at any strength level for an initiation codon variant. If there are no alternative start codons in the transcript set for a gene, then we recommend applying PVS1_Moderate for a start loss variant if one or more pathogenic variant(s) have been reported 5’ of the next downstream putative in-frame start codon (Methionine). On the other hand, if no pathogenic variant(s) occur upstream of the new Methionine then PVS1_Supporting should be applied.

##### Exonic deletions

The reading frame and NMD considerations illustrated for the ± 1,2 splice variants and the nonsense/frameshift variants (NMD only) are also applicable to single and multi-exon deletions. Whole gene deletions default to PVS1, assuming the gene in question meets the criterion for a LoF disease mechanism (**Table 1**). Although application of PVS1 (at a Very Strong level) would not reach a Pathogenic or Likely pathogenic classification using the combining rules in Richards et al 2017, the SVI working group acknowledged that for a full gene deletion of a known haploinsufficient gene, a Pathogenic classification is warranted as long as there is no conflicting evidence that would question the technical data or haploinsufficiency mechanism. Based on these considerations, we recommend interpreting the PVS1 rule for exonic variants as shown in **Figure 1**.

**TABLE 1.** Criteria for LoF disease mechanism.

|  |
| --- |
| • **Follow PVS1 Flowchart if:**<br>○ Clinical validity classification of gene is STRONG or DEFINITIVE<br>AND<br>○ 3 or more LOF variants are Pathogenic without PVS1 AND >10% of variants associated with the phenotype are LOF (must be across more than 1 exon) |
| • **Decrease final strength by <u>one</u> level (i.e. VeryStrong to Strong) if:**<br>○ Clinical validity classification of gene is at least MODERATE<br>AND<br>○ 2 or more LOF variants have been previously associated with the phenotype (must be across more than 1 exon)<br>AND<br>○ Null mouse model recapitulates disease phenotype |
| • **Decrease final strength by <u>two</u> levels (i.e. VeryStrong to Moderate) if:**<br>○ Clinical validity classification is at least MODERATE<br>AND EITHER<br>○ 2 or more LOF variants have been previously associated with the phenotype (must be across more than 1 exon)<br>OR<br>○ Null mouse model recapitulates disease phenotype |
| • **If there is no evidence that LOF variants cause disease, PVS1 should not be applied at any strength level.** |

##### Intra-genic duplications

In the clinical laboratory, duplications are most commonly identified through exon array, MLPA, CMA, or NGS-based algorithms. While the affected exon(s) may be readily identified using these technologies, the location of the duplicated region (i.e., intragenic or extragenic) is often unknown, which can in turn affect the pathogenicity of the variant. PVS1 should not be applied to exonic or whole gene duplications that are known to be inserted outside the relevant gene or if the duplication is a full gene inserted tandemly (**Figure 1**). If a duplication of a portion of the gene of a defined length is inserted in tandem, then one can predict if the reading frame will be disrupted leading to NMD, in which case PVS1 can be applied (**Figure 1**). PVS1 at any level should not be used if NMD is unlikely (or unknown) to occur since the underlying in-frame duplications of certain protein regions are not typically as disruptive as are their corresponding deletions.

Although one cannot assume duplications are in tandem, current data suggest that at least 83% of duplications (including exon level) are in tandem (Newman, Hermetz, Weckselblatt, & Rudd, 2015) (and unpublished data). Consequently, a duplication at an unknown insertion site is only downgraded one step to PVS1_Strong strength level provided it is predicted to shift the reading frame and cause NMD (**Figure 1**). Location and exact length of the duplicated fragment are essential to predict the affect on the protein’s reading frame. Uncertainty regarding a duplication length should preclude use of PVS1 given the inability to predict the affect on a protein’s reading frame and therefore NMD.

#### Disease Mechanism Considerations

PVS1 is only applicable if LoF is a disease mechanism for the relevant gene/disease association, as recommended in the ACMG/AMP guidelines (Richards et al). However, decsisions regarding use of the PVS1 strength level should also take into consideration the strength of evidence supporting the LoF disease mechanism for a given gene. For example, an LoF variant leading to a true null effect should have a stronger PVS1 level (e.g., PVS1_VeryStrong) if it affects a gene strongly linked to disease and wherein numerous pathogenic LoF variants have been reported compared to a gene with moderate evidence with only a limited number of LoF variants. A LoF variant might appropriately then have a lower (e.g., PVS1_Strong) evidence strength if it were present in the latter gene.

To provide guidance on how to weight a gene’s disease mechanism, we outline the general criteria as shown in **Table 1**. This is intended to provide a general framework until there is gene-level expert-curated mechanism information.

In general, the PVS1_VeryStrong pathogenic criterion can only be applied as shown in Figure 1 if used for a predicted LoF variant in genes with definitive or strong disease associations (Strande et al., 2017). Furthermore, we suggest it should only be applied if LoF variants make up at least 10% of the reported pathogenic variants in the gene and a minimum of 3 LoF variants have been classified as pathogenic without using the PVS1 rule. It is worth nothing that certain genes cause disease due to a LoF mechanism but do not harbor pathogenic LoF variants due to embryonic lethality and instead all pathogenic variants have milder impact such as leaky splice variants or mild missense variants. Therefore, while the above cutoffs might be inclusive for most definitive/strong gene-disease pairs with a LoF disease mechanism, we caution that some of those gene-disease pairs might not satisfy those cutoffs due to lethal LoF effects. Use of constraint scores as described below can be helpful in identifying genes of this type. On the other hand, no higher than PVS1_Strong should be assigned for a LoF variant in a gene with moderate disease evidence where only two LoF variants have been reported for a given phenotype and the phenotype is recapitulated by a knockout animal model. The evidence should be further downgraded to PVS1_Moderate for moderate genes meeting only one but not both of these criteria (**Table 1**).

For all the above criteria, the previously observed pathogenic LoF variants should be distributed across different exons of a given gene and the affected exon should not be alternatively spliced or lead to an in-frame effect.

Genes for diseases inherited in an autosomal dominant pattern should be particularly carefully assessed since disease mechanism (haploinsufficiency, gain of function, or dominant negative) is not necessarily established even for definitive or strong genes. Several resources, including the ClinGen haploinsufficiency (HI) score and the LoF constraint score (pLI or probability of LoF Intolerance) provided by the Exome Aggregation Consortium (ExAC), might be useful to assess if LoF is a potential disease mechanism for dominant genes. HI scores are divided into six tiers based on manually curated evidence (https://www.ncbi.nlm.nih.gov/projects/dbvar/clingen/). The pLI score measures the intolerance of a given gene to LoF variants in the general population such that a pLI > 0.9 [ref] suggests a significantly lower than expected rate of LoFs in this gene.

### VARIANT PILOT

Seven gene/disease-specific working groups (*CDH1*, *GAA*, *KCNQ1*, *PAH*, *PTEN*, *TP53*, and Hearing Loss) were tasked with testing the PVS1 flowchart each using five to ten LoF variants, of varying type, to determine if the evidence strength levels were appropriate for each variant (**Supplemental Table 1**). Of this pilot set, working groups agreed with the PVS1 evidence strength level for 89.3% (50/56) of variants. For the six discordant variants, working groups proposed a higher evidence level than specified in the PVS1 flowchart. For three of those six variants, working groups proposed a mechanism for elevating initiation codon variants with ≥1 pathogenic variant(s) upstream of the closest potential start codon from a Moderate strength to VeryStrong strength (PVS1). For example, variant NM_000152.4:c.1A>G in the *GAA* gene would reach PVS1_Moderate as the next in-frame methionine is at codon 122 of transcript NM_000152.4 and there are more than seven pathogenic or likely pathogenic variants in ClinVar between the methionine at codon 1 and the methionine at codon 122. Given the strength of evidence that variants lacking the region from codon 1 to codon 122 result in a nonfunctional protein product, the *GAA*/Pompe working group proposed that PVS1 be applied for initiation codons at its default strength level of very strong.

For the other three variants with discrepant strength levels between working groups and the PVS1 flowchart, disease groups proposed a mechanism for elevating variants that truncate or alter a region critical to protein function one in-frame exon deletion and two nonsense variants that escape NMD) from a Strong strength level to VeryStrong strength level (PVS1). For example, the *PTEN* working group applied PVS1 to variants that predicted a premature termination codon 5’ of the aspartic acid at codon 375 of transcript NM_000314.6. Codon 375 occurs in the middle of the last coding exon, meaning premature termination codons in the last exon 5’ of codon 375 are predicted to escape NMD and would have PVS1 applied at a Strong evidence level (PVS1_Strong) based on our proposed flowchart. However, truncation in the last *PTEN* exon upstream of codon 375 predicts the disruption of the C-terminal domain which includes PEST motifs, residues that undergo phosphorylation, and a PDZ domain-binding motif, which are critical to PTEN protein function. Although NMD is not predicted to occur, truncation of this region results in documented loss of function and thus the *PTEN* working group proposed PVS1 at a VeryStrong strength level be applied (Mester 2018; current issue).

After reviewing the PVS1 pilot variant results, the SVI working group elected to retain the current PVS1 evidence strength levels since our recommendations are meant to be a general guidance across all disease areas. The differences in classification represent the appropriate application of disease and gene-level specifications based on expert knowledge.

## CONCLUSION

The ACMG/AMP guidelines have been widely implemented by clinical laboratories and have been shown to promote consistent interpretations among laboratories; however, due to subjective interpretation of ACMG/AMP criteria, differences in their application still remain (Amendola et al., 2016; Harrison et al., 2017). ClinGen’s Sequence Variant Interpretation (SVI) Working Group, which has taken on the task of refining and evolving the current ACMG/AMP guideline to improve consistency in usage, has created recommendations for interpreting predicted LoF variants (PVS1 criterion). As this criterion is the only one assigned a VeryStrong evidence for pathogenicity in the original recommendations, caution is required to prevent overestimation of the variant impact and subsequent incorrect variant classification. The working group created a PVS1 decision tree to determine the appropriate strength level of PVS1 by addressing issues specific to each variant type (duplication, deletion, splice site, nonsense/frameshift, initiation codon) as well as recommendations for determining if LoF is a disease mechanism for the gene of interest. Correct usage of PVS1, with regard to variant impact and gene mechanism, will result in greater consistency in interpreting predicted LoF variants both from ClinGen Clinical Domain Working Groups and clinical laboratories. Future work will provide additional guidance regarding the combination of PVS1 with other rules.

## ACKNOWLEDGMENTS

Thank you to Jessica Mester (PTEN working group), Rachid Karam (CDH1 working group), Amanda Thomas (PAH working group), Catherine Rehder (GAA working group), and Melanie Care (KCNQ1 working group) for providing LOF variant examples. This research was supported in part by the National Human Genome Research Institute (NHGRI) under award U4100006834 and the Extramural and Intramural Research Programs of the National Human Genome Research Institute, National Institutes of Health. LGB was supported by the Intramural Research Program of the National Human Genome Research Institute grant number HG200359 09.

ClinGen Sequence Variant Work Group Members: Leslie G. Biesecker (co-chair), Steven M. Harrison (co-chair), Ahmad Abou Tayoun, Jonathan Berg, Steven E. Brenner, Fergus Couch, Garry Cutting, Sian Ellard, David Goldstein, Marc Greenblatt, Matt Hurles, Peter Kang, Izabela Karbassi, Rachel Karchin, Jessica Mester, Robert L. Nussbaum, Anne O’Donnell-Luria, Tina Pesaran, Sharon Plon, Heidi Rehm, Sean Tavtigian, and Scott Topper.

